# Low-input proteomics enables proteome and phosphoproteome-scale molecular phenotyping of separase-deficient oocytes

**DOI:** 10.64898/2026.06.17.732846

**Authors:** Sandra A. Touati, Veronique Legros, Jean Baptiste Boyer, Victor Cochard, Guillaume Chevreux, Katja Wassmann

## Abstract

We performed a comprehensive quantitative proteomic analysis of mouse oocytes using as few as 40 oocytes per condition, comparing wild-type and separase knockout oocytes at metaphase I and metaphase II. To this end, we generated a deep proteomic library spanning oocyte cell cycle stages, enabling the identification of numerous phosphosites without phosphopeptide enrichment. We further combined data-dependent (DDA) and data-independent (DIA) acquisition strategies, analyzed through multiple software pipelines in both library-based and library-free modes. Our results reveal extensive proteome remodeling during the metaphase I to metaphase II transition in wild-type oocytes, consistent with dynamic regulation of meiotic processes. As a proof of concept for our workflow, we asked whether separase knockout oocytes—unable to separate chromosomes in meiosis I—progress into meiosis II. Direct comparison of wild-type and separase knockout oocytes at the metaphase II stage revealed minimal global differences, supporting the idea that both conditions converge toward a comparable metaphase II-like cellular state despite distinct chromosomal configurations. However, at a finer scale, specific alterations were detected among chromosome-associated proteins. Notably, Meikin was enriched in separase-deficient metaphase II oocytes, consistent with defective separase-dependent cleavage and subsequent turnover. More broadly, several proteins involved in chromosome organization displayed behavior similar to Meikin, suggesting that separase activity regulates multiple substrates to orchestrate chromosome segregation during female meiosis.

## Introduction

Proteome and phosphoproteome analysis of synchronized cell cultures provide crucial insights into how mitotic kinases regulate cell cycle progression through substrate phosphorylation. This approach is particularly useful when cells can be synchronized and sufficient starting material is available. In a mutant context, these global mass spectrometry approaches allow the identification of downstream targets or pathways of specific kinases or phosphatases. In mammalian cells, this has led to the discovery of substrate consensus site preferences for key cell cycle phosphatases and kinases (Hein et al. 2017; Kruse et al. 2020; Hoermann et al. 2020; Touati 2026).

Meiosis, unlike mitosis, involves two divisions without an intermediate S-phase. Meiotic progression in oocytes and spermatocytes relies on tightly coordinated proteome remodeling and phosphorylation dynamics to ensure accurate chromosome segregation in two successive rounds: first, the segregation of homologous chromosomes (bivalents) during meiosis I, and second, the segregation of sister chromatids (dyads) during meiosis II (Petronczki et al. 2003; Marston and Amon 2004).

However, the scarcity of oocytes and the lack of synchrony in spermatozoa have limited global approaches to understanding how meiotic cell cycle progression is regulated in mammalian germ cells. In contrast, global mass spectrometry approaches in budding yeast have enabled the study of substrate preferences of key meiotic kinases, while alternative systems such as *Xenopus laevis* and starfish, which are evolutionarily closer to mammals, provide large quantities of oocytes for global analyses (Celebic et al. 2024; Koch et al. 2024; Wettstein et al. 2024). These systems have been used for global proteomics and phosphoproteomics studies. In starfish, a comprehensive study defined the phosphorylation landscape as oocytes undergo meiotic divisions and fertilization (Swartz et al. 2021), and Mos-MAPK targets were identified through phosphoproteomics (Avilov et al. 2025). However, unlike vertebrate oocytes, which are fertilized in meiosis II, starfish oocytes are fertilized in meiosis I, making them distinct and insufficient to study progression through meiosis I without comparison to a vertebrate model. In *Xenopus laevis*, previous studies compared several meiotic stages (Presler et al. 2017; Peshkin et al. 2025). However, the absence of a visible marker for meiosis I to meiosis II transition and the low temporal resolution from meiotic resumption to metaphase II, compared to mouse oocytes, limits the establishment of a time-resolved phosphorylation data set in meiosis I.

In mouse oocytes, a previous study identified 6700 proteins and 2600 phosphoproteins using replicates of 2000 oocytes each (Sun et al. 2024). The meiotic stages analyzed included prophase I (Germinal Vesicle (GV) or Germinal Vesicle BreakDown (GVBD), equivalent to nuclear envelope breakdown in mitosis), meiosis I (spanning GVBD until early prometaphase I), and metaphase II. Importantly, phosphopeptide enrichment was used to identify phosphorylated peptides, explaining why this study required such a high number of oocytes—a limitation when comparing different mutant mouse strains, where only a few mice are available.

Here, we aimed to develop a workflow for global mass spectrometry analysis using a significantly reduced number of mouse oocytes matured *in vitro*. Our goal was to achieve sufficient sensitivity from a minimal number of oocytes to perform this analysis at precise cell cycle stages and in a mutant context. Our study had three main objectives: (1) to establish a sensitive and robust workflow suitable for low-input samples, enabling the detection of key cell cycle markers across meiotic stages and providing a framework for determining oocyte cell cycle states in various mutants; (2) to precisely define the cell cycle state of oocytes lacking separase, a key protein required for chromosome segregation; and (3) to identify proteins and phosphoregulation events that behave differently in the presence or absence of separase.

Separase is a protease responsible for the cleavage of cohesin complexes and the resolution of chromosome cohesion. During mitosis, separase cleaves the kleisin subunit Rad21/Scc1, whereas in meiosis it primarily targets the meiotic kleisin Rec8, together with other substrates including Rad21L and the meiosis-specific kinetochore regulator Meikin, thereby coordinating homolog and sister chromatid segregation (Buonomo et al. 2000; Herrán et al. 2011; Maier et al. 2021; Yu et al. 2025). Intriguingly, in separase-deficient oocytes (here called KO oocytes), cell cycle progression appears to proceed beyond anaphase I despite the absence of cohesin cleavage, as illustrated by the degradation of securin (Kudo et al. 2006). However, how this uncoupling between cell cycle progression and chromosome segregation is reflected at the level of the proteome and phosphoproteome remains largely unknown.

Our study establishes a robust and sensitive low-input proteomic workflow, using as few as 40 oocytes (equivalent to the yield from 1–2 mice). This approach enables the identification of key cell cycle regulators in meiosis I and meiosis II, and can be broadly applied to determine the cell cycle stage and molecular state of oocytes in diverse mutants. The mass spectrometry workflow relies on data-independent acquisition (DIA) and integrates direct analysis with a deep spectral library, enabling the quantification of up to 6100 proteins per condition. We found that multiple cell cycle regulators—including BubR1, Cyclin B2, Mad2, Plk1, and Wee2—displayed changes in protein abundance between metaphase I and metaphase II, accompanied by specific regulation at the phosphosite level (Marston and Wassmann 2017). Remarkably, a similar global trend was observed in separase KO oocytes, indicating that large-scale proteome reorganization is mostly preserved in the absence of separase. Together, our findings demonstrate that our low-input proteomic workflow enables us to show that separase deficiency uncouples chromosome segregation from global cell cycle progression without substantially altering the overall proteomic landscape or key phosphosites at MII.

## Results

### A deep proteome library of mouse oocytes reveals extensive protein and PTM coverage without enrichment

To establish a comprehensive reference of protein and phosphorylation dynamics during meiotic progression, we generated a deep proteomic library from mouse oocytes undergoing meiotic maturation, across multiple cell cycle stages (Figure 1A). For *in vitro* maturation, oocytes can be obtained from adult female mice in GV stage corresponding to prophase I. Upon nuclear envelope breakdown (GVBD), they undergo a very long prometaphase and metaphase of meiosis I. Depending on the mouse strain, around 7-9 hours after GVBD oocytes undergo metaphase I to anaphase I transition. Oocytes proceed into meiosis II and establish a metaphase II arrest to await fertilization, around 12-14 hours after GVBD. Upon fertilization, anaphase II takes place, followed by exit from meiosis with the formation of a female pronucleus.

**Figure 1.**
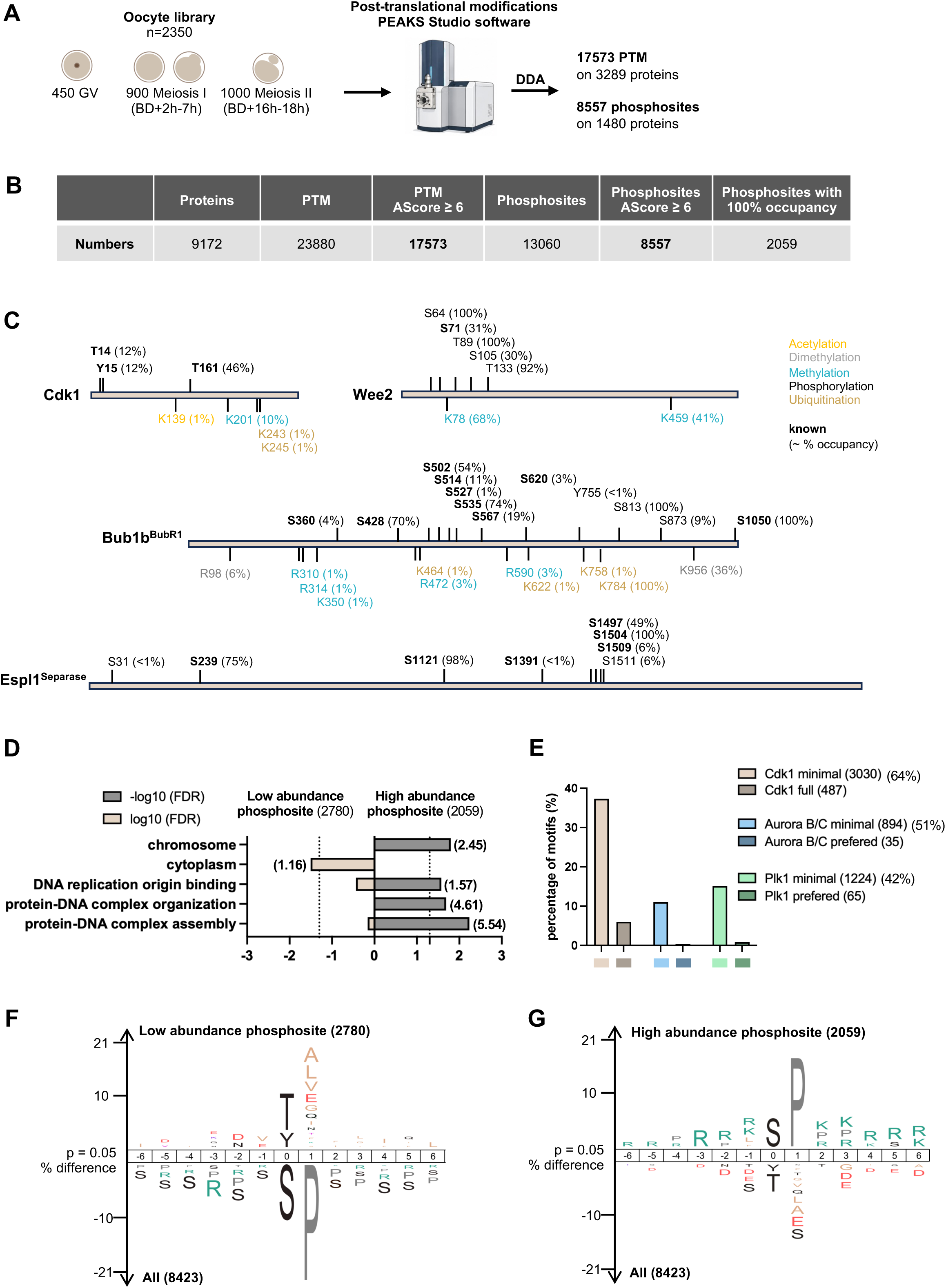
Comprehensive characterization of the oocyte proteome and phosphoproteome library. (A) Composition of the oocyte library, showing the number of oocytes at different meiotic stages. Across all samples, a total of 17523 PTM including 8423 phosphosites were quantified with high localization probability (AScore ≥ 6). (B) Summary table reporting the numbers of each category identified in the proteome and phosphoproteome. (C) Examples of proteins with quantified PTMs, indicating their position and percentage of phospho-abundance in brackets. Phosphorylations are indicated in black, with previously reported phosphosites shown in bold. The rest of PTM are indicated in their corresponding colors. (D) Gene Ontology enrichment analysis of proteins with high abundance phosphosites (grey, 100% abundance) ) compared with proteins with low abundance phosphosites (beige, <10% abundance). Bar length represents the log10(FDR) (beige, <10% abundance) or -log10(FDR) (grey, 100% abundance), and the value displayed in brackets next to each bar indicates the fold enrichment (only terms with significant enrichment are shown). Dashed line indicates the threshold p-value < 0.05. GO enrichment analysis was performed using the PANTHER classification system. GO terms cellular component (chromosome and cytoplasm), molecular function (DNA replication origin binding) and biological process (protein-DNA complex organization and assembly) are shown. (E) Distribution of key cell cycle kinase motifs across the phosphoproteome. bars the percentage of phosphosites within each motif and their phospho-abundance for every minimal motifs: Cdk1 minimal motif ((S/T)P), Cdk1 full motif ((S/T)Px(K/R)), Aurora B/C minimal motif ((R/K)x(S/T)), and Aurora B/C preferred motif ((R/K)(R/K)x(S/T)Φ), Plk1 minimal motif ((D/E)x(S/T)), Plk1 preferred motif ((D/E)x(S/T)Φx(D/E)). Minimal motifs define the core recognition sites of each kinase, while extended motifs capture sequence contexts that enhance phosphorylation efficiency and selectivity. (F-G) IceLogo motif analysis (p = 0.05) showing enrichment of sequence motifs. (F) Comparison between phosphosites with <10% abundance versus the remainder of the phosphoproteome. (G) Comparison between phosphosites with 100% abundance versus the remainder of the phosphoproteome.

In total, 450 GV oocytes, 900 oocytes in meiosis I (spanning prometaphase to anaphase I), and 1000 oocytes arrested in metaphase II were used to build the library. Notably, this corresponds to approximately 60 μg of total protein input. The sample was fractionated using high-pH chromatography (Batth et al. 2014) and fractions were analyzed using data-dependent acquisition (DDA) (see Materials and Methods). Across all samples, we identified 9172 proteins, corresponding to approximately 43% of the mouse protein-coding proteome. In addition, we identified a total of 17573 post-translational modifications (PTMs) with high localization confidence (AScore ≥ 6) (Figure 1A-B). Nearly half of the detected PTMs corresponded to phosphorylation events, with 8557 phosphosites identified, while the remaining modifications included acetylation, methylation, and ubiquitination (Figure S1A). The distribution of phosphorylated residues was consistent with expectations, with 74.6% serine, 21.9% threonine, and 3.5% tyrosine (Figure S1B). Importantly, this extensive coverage was achieved without any prior enrichment. Therefore, modified and unmodified peptide abundance can be used to estimate percentage of site occupancy. The number of phosphosites quantified in this study provides a good indication of the depth of analysis achievable with this workflow (Figure 1A-B).

We identified numerous examples of both previously reported phosphosites (highlighted in bold) and novel sites, underscoring the depth and quality of the dataset (Figure 1C). For instance, on Cdk1 (Cyclin Dependent kinase 1, master regulator of cell cycle progression), we detected the well-characterized inhibitory sites T14 and Y15, as well as the activating site T161 (Dupré and Wassmann 2025). In addition, multiple other PTMs, including acetylation, methylation, and ubiquitination, were identified on the same protein. We also identified several phosphosites on the kinase Wee2 (oocyte-specific Wee1 isoform that inhibits Cdk1 to control meiotic timing), including S71, previously reported by similarity using phosphopeptide enrichment strategies (Shimaoka et al. 2011). The protein BubR1 (gene *Bub1b*) (a key component of the spindle assembly checkpoint ensuring proper chromosome segregation) displayed a high density of PTMs including numerous newly identified sites (Elowe et al. 2007; Huang et al. 2008). Many of them cluster in its kinetochore localization or protein stability domains. Finally, we characterized multiple PTMs on separase (Espl1), including the Cdk1-dependent priming site S1121 and S1391, which are required for Cyclin B1 binding and separase inhibition, as well as previously described sites (S1497, S1504, S1509) located within the autocleavage region (Gorr et al. 2005; Hellmuth et al. 2015; Yu et al. 2023; Heinzle et al. 2025) (Figure 1C). These results highlight the remarkable depth of phosphosite identification achieved from limited starting material without enrichment. Importantly, the detection of numerous well-characterized sites alongside many previously unreported ones supports the robustness and reliability of the dataset (The UniProt Consortium et al. 2025).

Beyond site identification, our approach enabled estimation of site occupancy, revealing a broad spectrum ranging from low (<10%) to fully occupied (100%) sites (Figure S1C). For each PTM shown in Figure 1C, the estimated abundance is indicated in brackets. Notably, 2059 phosphosites (∼24% of all quantified sites) displayed abundance close to 100%, suggesting constitutive phosphorylation throughout meiotic progression, whereas the majority are likely dynamically regulated during the cell cycle. To assess the functional relevance of phosphorylation abundance, we performed Gene Ontology (GO) enrichment analyses comparing proteins harboring highly occupied (100%) versus lowly occupied (<10%) phosphosites (Figure 1D). This analysis revealed distinct functional enrichments: highly occupied phosphosites were preferentially associated with chromosome-related localization (GO term cellular component) and DNA replication origin binding (GO term molecular function), whereas low-abundance sites were enriched in cytoplasmic localisation and processes (Figure 1D). This pattern is consistent with the oocyte-specific meiotic program, in which DNA replication origins are maintained in an inhibited state through phosphorylation to prevent re-licensing between meiosis I and II and preserve genome integrity (Furuno et al. 1994; Lemaître et al. 2002; Daldello et al. 2015). We next examined kinase motif distribution, focusing on major cell cycle kinases including Cdk1, Aurora B/C, and Plk1. These kinases exhibit distinct substrate specificities that can be captured through minimal and preferred consensus motifs (Touati 2026). Motif analysis confirmed the widespread presence of these kinase signatures across the phosphoproteome, with both minimal and extended motifs detected (Figure 1E). Notably, Cdk1-associated sites displayed the highest overall abundance, whereas Plk1 motifs were generally associated with lower abundance. IceLogo motif analysis further supported these observations, revealing that phosphosites with low abundance (<10%) were enriched in threonine and tyrosine residues and in Plk1-like motifs (characterized by acidic residues such as aspartate or glutamate at the −2 position). In contrast, highly occupied sites (100%) were predominantly enriched in Cdk and Aurora consensus motifs, with Cdk motifs defined by a proline-directed sequence ((S/T)P) and Aurora motifs characterized by basic residues (arginine or lysine) in upstream positions, typically at −2 to −3 relative to the phosphosite (Figure 1F-G)(Kettenbach et al. 2011).More specifically, fully occupied Cdk sites showed strong enrichment for the extended Cdk consensus motif ((S/T)Px(K/R)), which includes a basic residue at the +3 position and is associated with increased substrate specificity and phosphorylation efficiency (note that Proline residues at the +1 position are downscaled in the sequence logos for improved visualization) (Figure S1D-E). Together, these results establish a high-resolution proteome and PTMs resource for mouse oocytes and demonstrate that extensive phosphorylation information, including site abundance and kinase signatures, can be robustly captured without phosphopeptide enrichment strategies.

The resulting dataset was then used to construct a spectral library, which serves as a reference for peptide identification and quantification in subsequent DIA experiments presented below. Analyses using this spectral library are hereafter referred to as « library-based approach », in contrast to « library-free » workflows. In downstream analyses, the spectral library was employed with Spectronaut (Biognosys, Zurich, Switzerland), which provides robust protein-level quantification in DIA, and with PEAKS Studio (Bioinformatics Solutions, Waterloo, Canada), which was rather used for PTMs analysis. The combined use of complementary software tools here was essential to maximize the extraction of meaningful biological information from the dataset (Figure S2A) (see Materials and Methods). This resource is publicly available on PRIDE and can be leveraged by the community for DIA-based proteomic analyses using Bruker Daltonics instrumentation.

### Spectral library based DIA improves proteome coverage in mouse oocytes

To assess whether our workflow can reliably capture proteome dynamics from minimal sample input in both physiological and mutant condition, we performed quantitative proteomic analyses in wild-type (WT) and Espl1^Separase^ ZP3-Cre conditional knockout (KO) oocytes (hereafter referred to as separase KO) (Kudo et al. 2006). Separase KO oocytes progress into metaphase II without chromosome segregation, judging from the degradation and reaccumulation of Securin. Metaphase II separase KO oocytes can be induced to undergo anaphase II with sister chromatid segregation, however, it was unknown whether oocytes without separase progress into a *bona fide* physiological metaphase II arrest for fertilization such as control oocytes (Gryaznova et al. 2021). To this end, we performed quantitative proteomic analyses using as few as 40 oocytes per condition (approximately 1 μg of total protein), comparing WT and separase KO at GVBD+6h (hereafter referred to as BD+6h and corresponding to metaphase I in WT) and GVBD+17h (hereafter referred to as BD+17h and corresponding to metaphase II arrest in WT) (Figure 2A). Data were acquired using DIA and analyzed with two complementary software pipelines (PEAKS Online and Studio and Spectronaut). In Spectronaut, both library-free and spectral library–based approaches were employed to improve protein identification, expand proteome coverage, and enhance the dynamic range of protein quantification (Figure 2A). A detailed overview of the data processing workflows for protein and PTMs identification is provided in Figure S2A.

**Figure 2.**
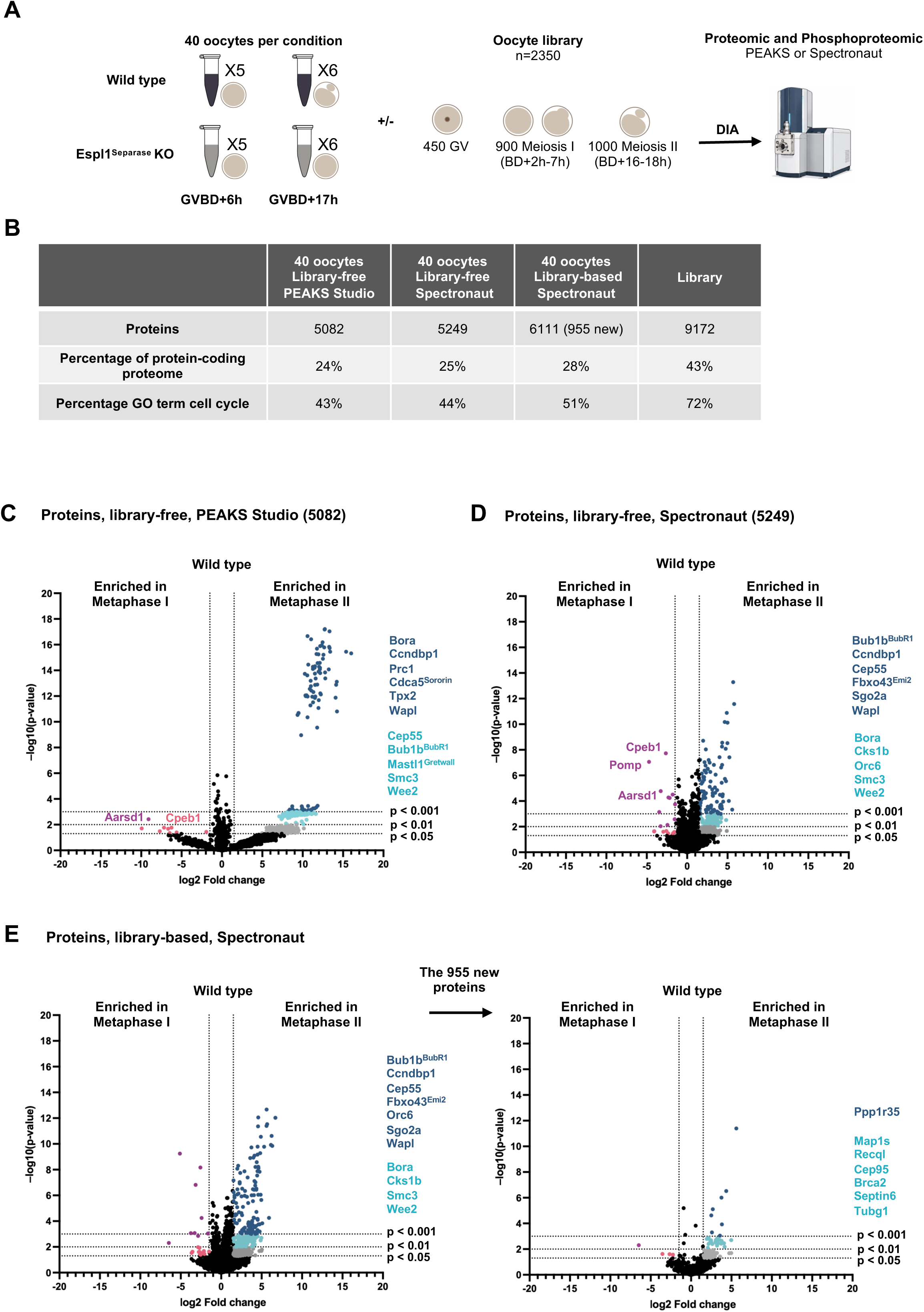
Global proteomic workflow and comparison of WT metaphase I and II oocytes across acquisition and analysis strategies. (A) Schematic overview of the experimental design. Proteomes were quantified from WT and Espl1^Separase^ KO mouse oocytes collected at metaphase I (GVBD+6h) and metaphase II (GVBD+17h). For each condition, the number of biological replicates and the number of oocytes per replicate are indicated. Data were processed using different software, with or without incorporation of the spectral library. (B) Summary table reporting the numbers of each category identified in the proteome and phosphoproteome. (C-E) Volcano plots for the pairwise comparison of WT metaphase I vs WT metaphase II. Dashed lines indicate the significance thresholds (fold-change ≥ 1.5 combined with either p < 0.05, p < 0.01 or p < 0.001, as indicated). Key cell cycle proteins enrichment is highlighted in colors depending on their p-value. Software used and, where applicable, the spectral library are indicated. Quantification of total proteins in each condition are shown in brackets. For panels (C) PEAKS Studio software (library-free), (D) Spectronaut software (library-free), and (E) Spectronaut software processed with the spectral library (left) with only newly identified proteins extracted (right).

Using either PEAKS Studio or Spectronaut in library-free mode, we quantified a similar number of proteins, corresponding to approximately 24–25% of the mouse protein-coding proteome and covering 43–44% of proteins annotated with the GO term “cell cycle”. Of note, the spectral library itself contains 72% of proteins associated with the GO term “cell cycle”. Importantly, combining DIA runs with fractionated raw data as Library Extension Runs in Spectronaut analysis enabled the identification of 955 additional proteins, increasing the total coverage to 28% of the coding proteome. This improvement primarily benefited low abundance proteins, including cell cycle regulators, increasing the representation of “cell cycle” GO term proteins to 51% (Figure 2B and S2B).

Overall, the use of a spectral library proved particularly effective in increasing the coverage of low abundance proteins, including key cell cycle regulators. Crucially, this library constitutes a reusable resource that can be leveraged for future analyses beyond the scope of this study. Notably, the same library-associated benefit for DIA protein quantification was not observed with PEAKS Studio.

### A robust low-input proteomic workflow captures meiotic proteome remodeling between metaphase I and metaphase II

First, we focused on comparing metaphase I and metaphase II in WT oocytes. Across all analysis strategies, we consistently detected extensive proteome remodeling between the two cell cycle stages (Figure 2C–E). Numerous cell cycle regulators, including Bora (an activator of Aurora A that promotes mitotic entry), BubR1, Cep55 (involved in cytokinesis), Cpeb1 (a regulator of mRNA translation during meiotic progression), Tpx2 (a microtubule-associated factor required for spindle assembly), and Wee2, showed consistent changes in abundance between the two stages, regardless of the analysis strategy used. Notably, the majority of these proteins (except Cpeb1) displayed increased abundance in metaphase II, supporting that meiosis II is predominantly associated with protein accumulation, likely driven by translational activation of stored maternal mRNAs (Christou-Kent et al. 2020). Proteins uniquely identified through the spectral library-based approach are highlighted in the right panel of Figure 2E. Several low abundance proteins, not detected in library-free analyses, such as subunits of PP1 (a major serine/threonine phosphatase regulating multiple cell cycle transitions), Brca2 (a key factor in homologous recombination and DNA repair), and Cep95 (a protein involved in regulation of microtubule organizing center) were successfully quantified in library-based analysis.

### Proteomic analysis reveals that separase-deficient oocytes reach a metaphase II–like state

We next aimed to determine the precise cell cycle state of separase KO oocytes and, more broadly, to assess whether our proteomic strategy can reliably distinguish meiotic stages across different conditions, including mutant conditions. Comparative proteomic analyses across four conditions (WT BD+6h, WT BD+17h, KO BD+6h, KO BD+17h), processing with Spectronaut and the Library Extension Runs, showed that the global proteomic changes observed between BD+6h and BD+17h in WT oocytes are largely recapitulated in separase KO oocytes. In both conditions, numerous proteins displayed similar stage-dependent changes in abundance between metaphase I and metaphase II (Figure 3A–B). In contrast, differences between WT and separase KO oocytes at BD+6h were more limited (Figure 3C). Strikingly, direct comparison between WT and separase KO oocytes at BD+17h revealed minimal differences at the global proteome level (Figure 3D). This trend was consistently observed across different processing strategies, including PEAKS Studio (Figure S3A–D) and Spectronaut in library-free mode (Figure S3E–H). Importantly, when considering only proteins newly identified through the Spectronaut library-based approach, the same overall patterns were observed (Figure S4A–D), demonstrating that the use of a spectral library reinforces the robustness of the analysis.

**Figure 3.**
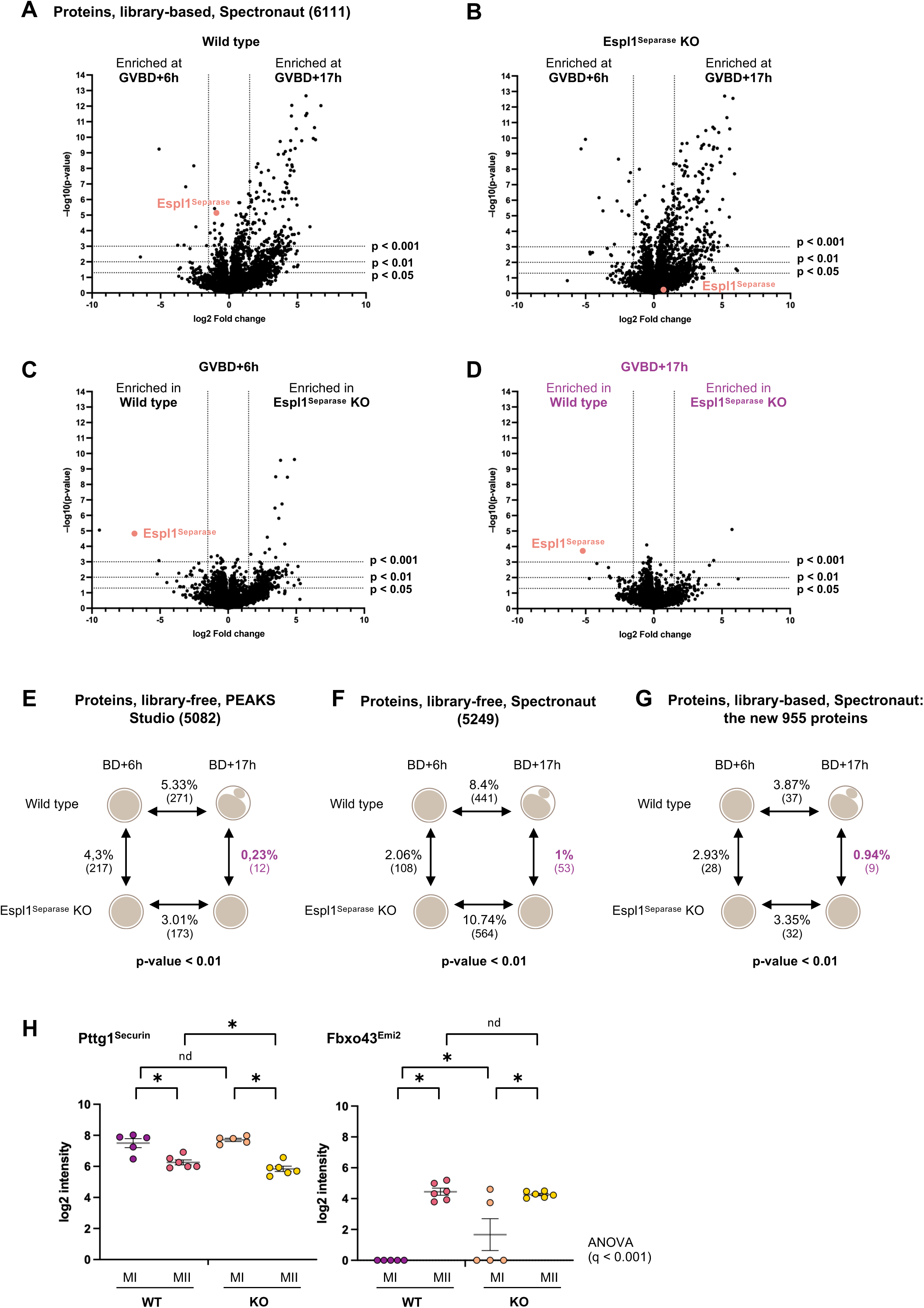
Proteomic evidence indicates that Espl1^Separase^ KO oocytes at GVBD+17h correspond to a metaphase II–like state. (A-D) Volcano plots for the four pairwise comparisons: WT GVBD+6h vs WT GVBD+17h, Espl1^Separase^ KO GVBD+6h vs Espl1^Separase^ KO GVBD+17h, WT vs Espl1^Separase^ KO at GVBD+6h, and WT vs Espl1^Separase^ KO at GVBD+17h. Dashed lines indicate the significance thresholds (fold-change ≥ 1.5 combined with either p < 0.05, p < 0.01 or p < 0.001, as indicated). The Espl1^Separase^ protein is highlighted in red in all plots and the WT and Espl1^Separase^ KO mouse at GVBD+17h is highlighted in purple. (E-G) Summary schematic indicating the percentage and numbers (indicated in parentheses) of differing proteins between each pair of conditions based on fold-change ≥ 1.5 and p < 0.01. The percentage difference for WT and Espl1^Separase^ KO mouse at GVBD+17h is highlighted in purple. Software used, and, where applicable, the spectral library are indicated. Quantifications of total proteins in each condition are shown in parentheses. (H) Plots of log2-transformed protein intensities for four conditions: WT MI, WT MII, KO MI, and KO MII (n = 5-6 biological replicates per condition). Each point represents a single protein measurement. One-way ANOVA was used to assess global variation (q < 0.001 as indicated), followed by FDR-corrected pairwise post-hoc t-tests. Statistical significance after FDR correction (0.05) is indicated by asterisks *. ‘nd’ indicates proteins that did not reach significance. Asterisks denote “discovery”, not the exact q value.” Boxplot indicates Mean with SEM.

A quantitative comparison of the proportion of significantly changing proteins across conditions further supported these observations. The largest differences were consistently observed between BD+6h and BD+17h, in both WT and separase KO oocytes (ranging from 3% to 10.74%, depending on the software and analysis strategy) (Figure 3E–G). In contrast, fewer changes were detected between WT and separase KO oocytes at BD+6h (2% to 4.3%), and only minimal differences were observed at BD+17h (0.23% to 1%) (Figure 3E–G). Together, these results strongly suggest that separase KO oocytes converge toward a metaphase II–like proteomic state as WT oocytes.

To further validate this conclusion at the level of individual proteins, we examined well-characterized cell cycle markers. Securin (Pttg1), a known inhibitor of separase, is degraded at the metaphase I-anaphase I transition and subsequently reaccumulates at lower levels during metaphase II. Conversely, the protein Emi2 (*Fbxo43* gene) accumulates during meiosis II and is required to maintain metaphase II arrest (Bouftas et al. 2022). Both proteins displayed similar abundance dynamics in WT and separase KO oocytes (Figure 3H), further arguing that separase KO oocytes are arrested in a metaphase II–like state.

Together, these results provide strong proteomic evidence that separase-deficient oocytes progress to a metaphase II–like state despite defective chromosome segregation. More broadly, this low-input proteomic strategy enables precise determination of the cell cycle state of oocytes and can be applied to characterize diverse mutant conditions.

### Proteomic identification of conserved metaphase I and metaphase II markers in wild-type and separase-deficient oocytes

To further validate cell cycle staging at the protein level and get a better understanding of proteins with more abundance in metaphase I or metaphase II, we examined in detail proteins differentially abundant between these meiotic stages in both WT and separase KO oocytes. Initial pairwise comparisons (t-tests) in WT oocytes already revealed stage-specific differences, with a strong enrichment of proteins more abundant in metaphase II (Figure 2). These “increasing” markers are particularly informative as they reflect the progressive accumulation of proteins required for meiosis II and metaphase II arrest. To obtain a more global and comparative view across all conditions, we performed a one-way ANOVA across all four conditions (WT MI, WT MII, KO MI, KO MII).

This analysis confirmed that several proteins identified as enriched in metaphase II in WT oocytes including BubR1, Ccndbp1 (Cyclin D–binding protein 1, involved in cyclin-dependent regulation of the cell cycle), Tpx2, and Wee2 follow the same trend in separase-deficient oocytes (Figure 4A-S4E). Importantly, these increasing markers behave consistently across WT and mutant conditions, making them robust indicators of progression toward a metaphase II state.

**Figure 4.**
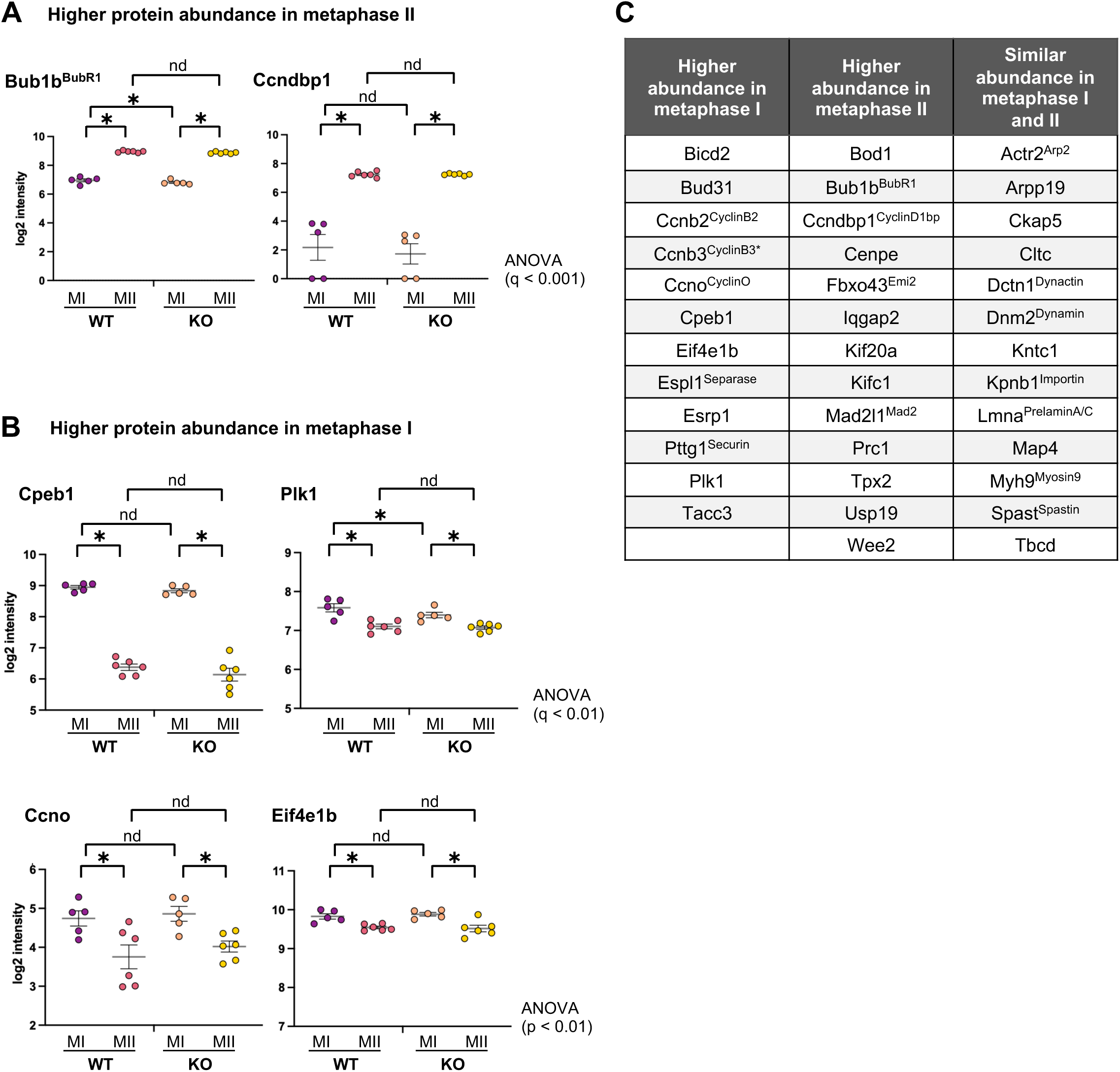
Identification of new protein markers distinguishing metaphase I and metaphase II oocytes. (A-B) Plots of log2-transformed protein intensities for four conditions: WT MI, WT MII, KO MI, and KO MII (n = 5-6 biological replicates per condition). Each point represents a single protein measurement. One-way ANOVA was used to assess global variation (q or p-value is indicated directly on the figure), followed by FDR-corrected pairwise post-hoc t-tests. Statistical significance after FDR correction (0.05) is indicated by asterisks *. ‘nd’ indicates proteins that did not reach significance. Asterisks denote “discovery”, not the exact q value. Boxplot indicates Mean with SEM. (A) Proteins more abundant in metaphase II. (B) Proteins more abundant in metaphase I. C) Proteins are grouped as metaphase I-enriched, metaphase II-enriched, or stable based on their relative abundance. Protein names are shown. Markers are consistently identified across datasets and provide a robust framework to discriminate meiotic stages. Cyclin B3 is included based on prior work but remains of low abundance in this dataset, limiting confident quantification, and is shown by an asterisk.

ANOVA analysis further revealed cell cycle proteins enriched in metaphase I. These MI-enriched proteins, including Plk1 (Polo-like kinase 1, regulating multiple steps of mitotic and meiotic progression), Ccno (Cyclin O, involved in cell cycle regulation), and Eif4e1b (a translation initiation factor), displayed consistent abundance patterns in both WT and separase-deficient oocytes (Figure 4B). These “decreasing” markers are equally important, providing complementary information to increasing markers. Figure 4C provides a comprehensive summary of these protein markers, including proteins enriched in metaphase I, proteins enriched in metaphase II, and proteins displaying stable abundance across stages. Notably, Cyclin B3, previously identified in our laboratory as a key regulator of meiotic progression and known to be absent at metaphase II (Karasu et al. 2019; Bouftas et al. 2022; Dupré and Wassmann 2025), is also included in this framework; however, its low abundance in our dataset limits robust quantification.

Together, these complementary analyses highlight the importance of combining pairwise and multi-group statistical approaches to capture the full spectrum of proteome dynamics. They also define a robust set of protein markers distinguishing metaphase I and metaphase II, applicable across both WT and mutant conditions. The combination of increasing (MII-enriched) and decreasing (MI-enriched) markers provides a bidirectional framework to define cell cycle progression. This is particularly critical in a mutant context, where specific pathways may selectively affect protein turnover, *i.e* accumulation or degradation. In such cases, relying on only one class of markers could lead to misinterpretation of the cell cycle state. Importantly, these markers confirm that separase-deficient oocytes at BD+17h exhibit a protein expression profile characteristic of metaphase II, reinforcing the conclusion that global cell cycle progression is largely preserved despite defective chromosome segregation (Kudo et al. 2006).

### The phosphoproteome of separase-deficient oocytes is globally preserved at metaphase II

We next investigated if the phosphorylation state of separase KO oocytes confirms progression into metaphase II, by analyzing general post-translational modifications (PTMs) and more precisely phosphopeptides. We used the peptide-centric DIA data acquired and searched them against the comprehensive and high quality spectral-library generated from DDA experiments after extensive fractionation. This approach is gaining popularity, especially because it allows users to investigate PTMs by taking advantage of the systematic and unbiased recording of fragmentation information from DIA data to leverage better quantification results (Yang and Qiao 2023). Here, we investigate for the first time these tools in the context of the cell cycle progression in mouse oocytes in order to estimate its potential to extract meaningful PTMs data with challenging amounts of starting material.

4639 modified peptides including phosphorylation, ubiquitination, methylation and acetylation were identified with a similar distribution as in the oocyte library. A detailed classification of different PTM categories is provided in Figure S5A. Global PTM and phosphoproteomic analyses revealed that, similarly to the proteome, the post-translational modification landscape and phosphoproteome of separase KO oocytes at metaphase II closely resemble those of WT MII oocytes. Among the 4639 peptides carrying at least one PTM, only 0.45% (highlighted in purple) showed significant differences between WT and KO oocytes at this stage, whereas substantially larger changes were observed between metaphase I and metaphase II in both conditions (2.28% and 2.34%, respectively) (Figure 5A). More specifically, among the 1136 phosphopeptides carrying a single phosphorylation site, only 0.44% (highlighted in purple) differed significantly between WT and KO at this stage, compared to markedly higher proportions observed between metaphase I and metaphase II in WT and KO oocytes (4.84%) (Figure 5B and S5B–E).

**Figure 5.**
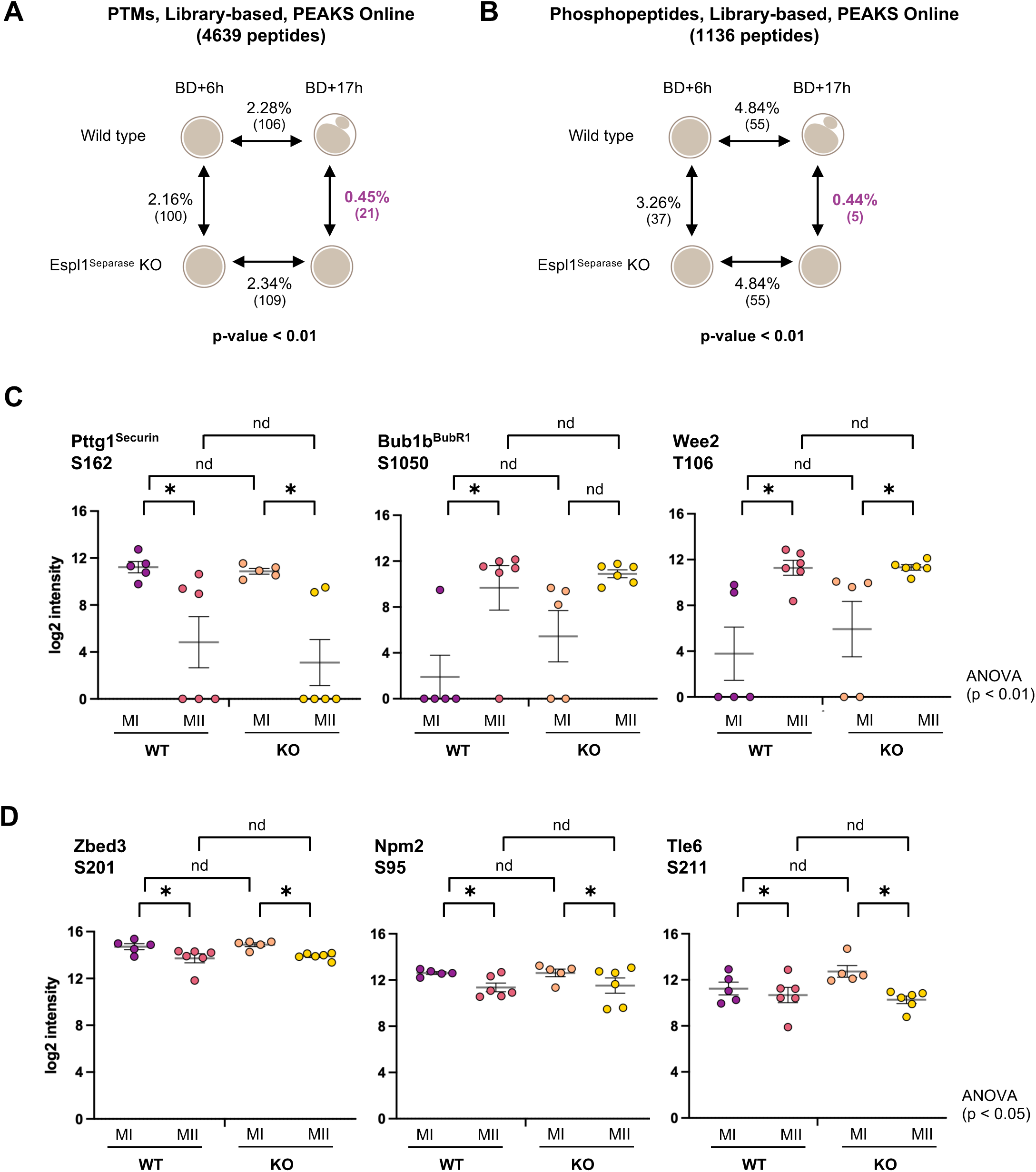
Phosphoproteomic evidence indicates that Espl1^Separase^ KO oocytes at GVBD+17h correspond to a metaphase II–like state. (A-B) Summary schematic indicating the percentage and numbers (indicated in brackets) of differing PTMpeptides (A) or phosphopeptides (B) between each pair of conditions based on fold-change ≥ 1.5 and p < 0.01. The percentage difference for WT and Espl1^Separase^ KO mouse at GVBD+17h is highlighted in purple. PEAKS Online software with the spectral library was used. Quantifications of total proteins in each condition are shown in parentheses. (C-D) Plots of log2-transformed protein intensities for four conditions: WT MI, WT MII, KO MI, and KO MII (n = 5-6 biological replicates per condition). Each point represents a single protein measurement. One-way ANOVA was used to assess global variation (q or p-value is indicated directly on the figure), followed by FDR-corrected pairwise post-hoc t-tests. Statistical significance after FDR correction (0.05) is indicated by asterisks *. ‘nd’ indicates proteins that did not reach significance. Asterisks denote “discovery”, not the exact q value. Note that due to the high number of zero values “0”, boxplots are not shown.

Among the 1136 phosphopeptides quantified, representative examples highlighted proteins exhibiting coordinated changes at both the proteome and phosphoproteome levels (PTMs and protein abundances quantification from the PEAKS software suite were only compared together). For instance, the Securin S162 phosphosite decreased between metaphase I and metaphase II in both WT and KO oocytes, consistent with the reduction in total Securin protein levels. Similarly, Bub1b^BubR1^ S1050 and Wee2 T106 showed increased phosphorylation in metaphase II, in line with their higher protein abundance at this stage (Figure 4C, 5C). In contrast, another category of phosphosites displayed stable protein abundance but dynamic phosphorylation (Figure 5D and S5F). This was the case for Zbed3 S201 (a zinc finger protein), Npm2 S95 (a nucleoplasmin protein), and Tle6 S211 (a translucin like enhancer protein) (Figure 5D). These examples highlight phosphorylation events that are regulated independently of total protein levels. Together, these findings indicate that the global phosphorylation programs associated with meiotic progression are largely maintained in the absence of separase.

### Separase deficiency reveals selective alterations in chromosome-associated proteins in metaphase II

Despite the overall similarity between WT and separase KO oocytes at the global proteome level, we next sought to identify more specific protein alterations that could reflect separase activity, particularly among chromosome-associated factors. Given that only a limited number of direct separase substrates have been described, we reasoned that our analysis might uncover additional candidates or reveal indirect effects of separase deficiency on chromosomal protein composition. Rec8 is the canonical meiotic kleisin subunit replacing the mitotic cohesin subunit Rad21, while Rad21L is also a meiotic kleisin but is present at much lower abundance than Rec8 in mouse oocytes (Lee and Hirano 2011). Notably, Rad21, was enriched in WT oocytes compared to separase KO oocytes. This protein, detected at low abundance and only through the Spectronaut library-based approach, displayed an increase in abundance between metaphase I and metaphase II in WT oocytes that was not observed in separase-deficient oocytes (Figure 6A–B). Given that Rec8 is the main meiotic kleisin responsible for cohesion in oocytes, and that it is normally replaced by Rad21 during mitotic divisions, one possibility is that Rec8 cleavage on chromosome arms during the MI–MII transition in WT oocytes may be required to enable Rad21 accumulation in meiosis II. The observed difference in Rad21 abundance between WT and separase KO MII oocytes may therefore reflect the persistence of bivalents associated to Rec8, which could indirectly alter Rad21 protein dynamics. Note, that unfortunately, we cannot detect Rec8 in our datasets.

**Figure 6.**
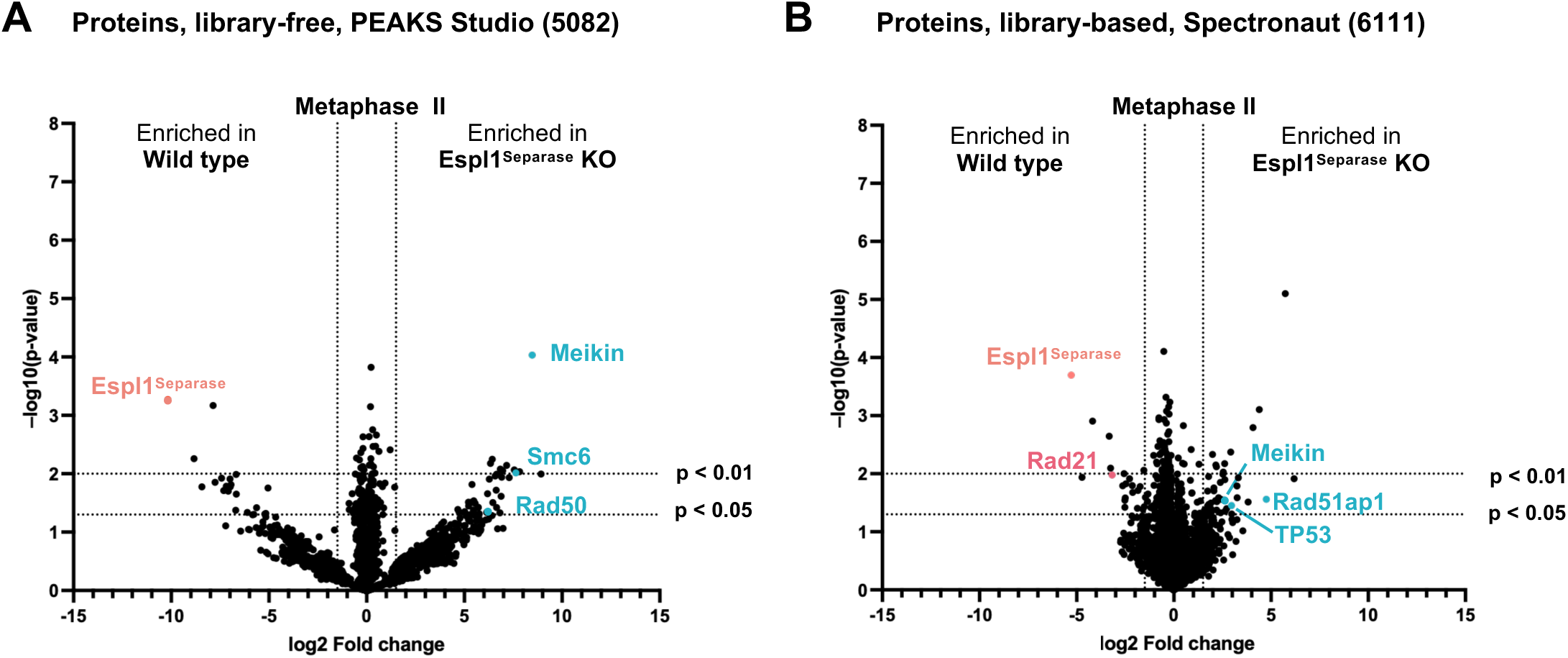
Chromosome-associated proteins differentially abundant in Espl1Separase KO metaphase II oocytes. (A-B) Volcano plot comparing WT vs Espl1^Separase^ KO oocytes in metaphase II. Dashed lines indicate significance thresholds (fold-change ≥ 1.5 combined with p < 0.05 or p < 0.01). Espl1^Separase^ protein is highlighted in red, the more abundant proteins in Espl1^Separase^ KO oocytes in blue and the less abundant in pink. (A) Results from PEAKS Studio software. (B) Results from Spectronaut software coupled with the spectral library.

In contrast, the meiosis-specific kinetochore regulator Meikin, a known separase substrate (Maier et al. 2021), was consistently enriched in separase KO oocytes (Figure 6A–B). This observation is fully consistent with defective separase-dependent cleavage and subsequent turnover. More broadly, this suggests that separase-mediated cleavage may contribute to the destabilization or degradation of specific chromosome-associated proteins, explaining their increased abundance in the absence of separase activity. Extending this analysis, several additional chromosome-associated proteins, including Rad50 (involved in DNA double-strand break repair), Smc6 (a component of the SMC5/6 complex important for chromosome maintenance), Rad51ap1 (a regulator of homologous recombination), and p53 (TP53, a key regulator of DNA damage responses), followed a similar trend to Meikin and were enriched in separase KO metaphase II oocytes (Figure 6A–B).

Together, these results indicate that, while the global proteome and phosphoproteome remain largely unchanged, separase deficiency leads to selective alterations in chromosome-associated proteins. These changes likely reflect a combination of direct effects on separase substrates and indirect consequences of altered chromosome organization.

## Discussion

This study establishes a robust and sensitive low-input proteomic framework enabling quantitative analysis at both protein and PTM levels from as few as 40 mouse oocytes. By combining a deep spectral library with DIA-based acquisition and analysis strategies, we demonstrate that oocyte cell cycle stages can be precisely resolved and molecular alterations reliably detected even in mutant conditions. This integrative workflow substantially improves the depth of the quantified proteome that is particularly useful to monitor low abundance regulatory proteins, while also providing a reusable resource for future studies. These advances are especially important in the context of mammalian oocytes, where proteomic analyses have historically been constrained by the scarcity of biological material and where phosphoproteomics has remained technically challenging.

Beyond the technical advance, our dataset provides insight into the molecular remodeling occurring during meiotic progression. A key feature emerging from the analysis is the identification of complementary classes of protein markers that either increase or decrease between metaphase I and metaphase II. These groups likely reflect distinct regulatory mechanisms and should be considered together to accurately define meiotic progression. In mammalian oocytes, progression through meiosis relies heavily on the precise temporal control of maternal mRNA translation, which counterbalances protein degradation during the MI-to-MII transition to enable entry into meiosis II (Susor and Kubelka 2017; Meneau et al. 2020). Consistent with this, our data reveal a marked accumulation of proteins between metaphase I and metaphase II, reflecting active translation from stored maternal mRNAs. This process is driven not by a global increase in translation, but rather by selective activation of specific mRNA subsets. Key regulators such as CPEB1 function as molecular switches by recruiting the translational machinery to target mRNAs through cytoplasmic polyadenylation elements (Radford et al. 2008; Huang et al. 2023). Interestingly, we observe that CPEB1 abundance decreases during meiosis II after polyadenylation has been initiated, while translation of already activated mRNAs is maintained. Together, these observations illustrate the highly stage-specific and tightly regulated nature of translational control during oocyte maturation, enabling the accumulation of proteins required for metaphase II progression while selectively turning off regulatory factors once their role has been fulfilled. In contrast, proteins decreasing in abundance between metaphase I and metaphase II represented a comparatively smaller group, supporting that large-scale protein degradation is not the dominant driver of proteome remodeling during meiosis I and II (Jentoft and Schuh 2025). A subset of these proteins may reflect canonical APC/C-dependent degradation pathways associated with meiotic progression (Pesin and Orr-Weaver 2008). However, additional mechanisms are also likely to contribute, including selective translational repression or unequal partitioning during polar body extrusion.

A major finding of this work is that separase-deficient oocytes, despite retaining bivalent chromosomes due to defective cohesin cleavage, nonetheless undergo extensive proteome and phosphoproteome remodeling and acquire a molecular state highly similar to metaphase II oocytes. This demonstrates that global cell cycle progression, as reflected by both protein abundance and phosphorylation dynamics, can proceed largely independently of chromosome segregation. These findings reveal a striking uncoupling between chromosome morphology and cell cycle progression in mammalian oocytes They highlight the robustness of the meiotic maturation program to generate an egg that can be fertilized, at the molecular level.

At the global scale, both the proteome and phosphoproteome displayed highly similar dynamics between wild-type and separase-deficient oocytes, with only limited differences detected at metaphase II. Furthermore, the identification of stage-specific protein markers and phosphorylation patterns confirmed that separase-deficient oocytes adopt a molecular signature characteristic of metaphase II despite their abnormal chromosomal configuration.

Although global remodeling was remarkably preserved, localized chromosome-associated alterations remained detectable. At a finer scale, our analysis uncovered selective enrichment of specific chromosome-associated proteins in separase-deficient oocytes. The accumulation of Meikin is fully consistent with its known cleavage by separase and supports a model in which separase activity contributes to the turnover of specific meiotic regulators (Maier et al. 2021). Importantly, despite persistent Meikin accumulation, separase-deficient oocytes still acquire a molecular signature characteristic of metaphase II, indicating that Meikin cleavage is not strictly required for progression toward a metaphase II–like state. Maintenance of meiosis I chromosome architecture and establishment of metaphase II molecular identity can therefore become uncoupled. More broadly, the enrichment of additional chromosome-associated factors, including proteins involved in DNA repair and chromosome maintenance, suggests that separase activity may directly or indirectly influence the composition, accessibility, or stability of chromosomal protein complexes.

Beyond the biological insights obtained here, this work highlights the broader potential of low-input proteomics for resolving cell cycle states in rare cell populations. The ability to accurately stage oocytes based on combined proteomic and phosphoproteomic signatures opens new opportunities for the analysis of meiotic mutants. Importantly, the combined use of increasing and decreasing protein markers provides a more robust and nuanced readout of cell cycle progression, particularly in situations where specific regulatory pathways are selectively impaired. More broadly, this strategy offers a framework for distinguishing defects in chromosome architecture, translation, degradation, or conditions in which these processes become uncoupled. In principle, this approach could be extended to additional meiotic mutants, including Cyclin B3-deficient oocytes, which were shown to arrest in a CSF-like metaphase I state (Bouftas et al. 2022). In such a context, translation-dependent accumulation of proteins may still occur, whereas proteins normally subjected to degradation or inactivation may remain abnormally stable. Proteomic profiling therefore provides an unbiased approach to determine the precise molecular state of mutant oocytes and to dissect the mechanisms underlying meiotic arrest phenotypes.

Importantly, although substantial phosphoproteome coverage was achieved without phosphopeptide enrichment, further improvements remain possible to increase the depth of analysis and enable the study of PTM dynamics in additional mutant conditions. In particular, expanding the spectral library using phospho-enriched datasets, as well as leveraging more sensitive instrumentation developed for single-cell analysis in combination with small-scale fractionation, are expected to further enhance coverage. Overall, these advances indicate that mass spectrometry has reached a level of maturity that now enables the investigation of cell cycle regulation processes that were previously limited by sensitivity constraints. Such approaches will be especially valuable for the future analysis of kinase and phosphatase mutants, where phosphorylation dynamics are central to the phenotype.

Altogether, this work establishes a framework for integrating proteomic and phosphoproteomic analyses into the study of meiotic regulation and mutant phenotypes in mammalian oocytes. By overcoming longstanding technical limitations, it opens the way for systematic and high-resolution dissection of the molecular logic governing meiotic progression. More broadly, our findings demonstrate that proteomic identity and chromosome morphology can become uncoupled during meiosis, revealing an unexpected robustness of the meiotic program despite severe defects in chromosome segregation.

## Materials and Methods

### Animal housing and ethical statement

Mice were maintained in an enriched environment with *ad libitum* access to food and water, in the conventional animal facilities of UMR7592 (authorization C75-13-17), following current French regulations. Housing conditions included a 12-h light/12-h dark cycle, in accordance with FELASA recommendations. All experiments were reviewed and approved by the French Ministry of Higher Education and Research (authorization n° B-75-0513), following national guidelines and the principles of the 3Rs to minimize animal use (Licence 5330). The number of animals used was kept to the minimum necessary to achieve statistically meaningful results. Female mice were either bred in-house at UMR7592 (including genetic knockouts and littermate controls) or purchased at 7 weeks of age (C57BL/6-Jrj or Swiss, Janvier Labs, France) and maintained until full sexual maturity. For Espl1^separase^ KO experiments, the mice described in Kudo et al., 2006 were used (Kudo et al. 2006). For all procedures, mice were euthanized by cervical dislocation between 9 and 12 weeks of age by trained personnel. Except for genotyping, no experimental manipulation was performed prior to euthanasia.

### Mouse oocyte collection and culture

Fully grown germinal vesicle (GV) stage oocytes were collected from ovaries of sexually mature female mice (9–12 weeks old). Ovaries were dissected in pre-warmed, self-prepared M2 medium supplemented with 100 µg/mL dbcAMP (dibutyryl cyclic AMP, Sigma-Aldrich, D0260) and covered with mineral oil (embryo culture grade, FUJIFILM Irvine Scientific, 9305) to maintain prophase I arrest. For *in vitro* meiotic resumption, oocytes were extensively washed to remove dbcAMP and transferred into dbcAMP-free medium under mineral oil, then cultured at 38°C (El Jailani et al. 2024; Cladière et al. 2025). Germinal vesicle breakdown (GVBD) was monitored and used as the temporal reference point (defined as time 0). Oocytes were collected at defined time points corresponding to specific meiotic stages: GVBD + 6 h (BD+6h), corresponding to metaphase I in wild-type oocytes, and GVBD + 17 h (BD+17h), corresponding to metaphase II arrest. For separase-deficient conditions, oocytes were obtained from Espl1^Separase^ ZP3-Cre conditional knockout females, also referred to as “separase KO” (Kudo et al. 2006). Oocytes from wild-type littermates were used as controls and processed in parallel under identical conditions. At each time point, oocytes were collected, washed in PBS to remove residual medium, and snap-frozen in minimal volume. Groups of 40 oocytes per biological replicate were used for downstream proteomic analyses. Separate groups of wild-type Swiss oocytes were pooled to generate the spectral library.

### Materials for MS

MS grade Acetonitrile (ACN), MS grade H_2_O, MS grade formic acid (FA), Tris(2-Carboxyethyl) Phosphine Hydrochloride (TCEP-HCl) and S-Methyl methanethiosulphonate (MMTS) were from ThermoFisher Scientific (Waltham, MA, USA). Sequencing-grade Trypsin/Lys C mix was from Promega (Madison, WI, USA). Ammonium bicarbonate (NH_4_HCO_3_), urea and ammonium formate were from Sigma-Aldrich (Saint-Louis, MO, USA).

### Preparation of the “low-input” samples

Samples (40 oocytes) were resuspended in 20µL of a denaturing buffer composed of 8 M urea-25 mM NH₄HCO₃ pH 7.8. They were reduced with 10 mM TCEP-HCl and alkylated with 20 mM MMTS. After a 16-fold dilution in NH₄HCO₃, proteins were digested overnight at 37 °C using a Trypsin/Lys-C mix at a 1:50 enzyme-to-substrate ratio. Prior to LC–MS/MS analysis, peptides were loaded onto Evotips (Evosep, Odense, Denmark) and desalted according to the manufacturer’s instructions.

### Preparation of the “oocyte library” samples

About 2000 mouse oocytes (50 µg) were digested as described previously. The peptide mixture was then acidified to a final concentration of 1% trifluoroacetic acid (TFA) and desalted using a Sep-Pak cartridge (Waters, Milford, MA, USA) following the manufacturer recommendation. The desalted peptides were dried under vacuum and resuspended in 20 mM ammonium formate buffer (pH 10). 20µg of peptides were injected onto a CSH Premier C18 column (2.1 x150 mm, 130Å, 1.7µm beads, Waters) equilibrated at 40°C and operated at a flow rate of 200 µL/min using an Acquity Premier UPLC fitted with a fraction collector (Waters, Milford, MA, USA). H_2_O-20mM NH_4_HCO_2_ and ACN/H_2_O-20mM NH_4_HCO_2_ (80/20) at pH 10 were used as solvent A and B respectively. Peptides were separated using a linear gradient of B from 0 to 50% over 50-min. A total of 24 fractions were collected every 2-min, dried under vacuum and resuspended for loading onto Evotips.

### LC-MS/MS acquisition

Samples were analyzed on a timsTOF Pro HT mass spectrometer (Bruker Daltonics, Bremen, Germany) coupled to an Evosep one system (Evosep, Odense, Denmark) operating with the 40SPD method developed by the manufacturer. Briefly, the method is based on a 32-min gradient and a total cycle time of 35 min with a C18 analytical column (0.075 x 150 mm, 1.9µm beads, IonOptocks) equilibrated at 50°C and operated at a flow rate of 200 nL/min. H_2_O-0.1 % FA was used as solvent A and ACN-0.1 % FA as solvent B. “Low input” samples were analyzed with a DIA-PASEF method comprising 12 pydiAID frames with 3 mass windows per frame resulting in a cycle time of 0.975 seconds as described in Bruker application note LCMS 218. “Oocyte library” fractions were acquired in DDA-PASEF mode with an accumulation time of 150 ms, 4 PASEF scans per cycle, covering an ion mobility range of 0.75–1.25 Vs/cm² and resulting in a cycle time of 0.78 s.

### Data analysis

#### Protein search

DIA raw files were processed with Spectronaut (version 20.5.260227.92449) using a direct DIA set up first and secondly using fractionated raw data as Library Extension Runs. Data were searched against the SwissProt Mus musculus database (04 2025, 17222 entries). Specific tryptic cleavages were selected and a maximum of 2 missed cleavages were allowed. The following post-translational modifications were considered for identification: Acetyl (Protein N-term), Oxidation (M), Deamidation (NQ) as variable and Beta-methylthiolation (C) as fixed. The maximum number of variable modifications was set to 3. Identifications were filtered based on a 1% precursor and protein Qvalue cutoff threshold. The protein LFQ method was set to automatic and the quantity was set at the MS2 level with a cross-run normalization applied. PEAKS Studio 13.1 was also used to process DIA raw files in a direct DIA workflow using same protein database and modification parameters. Identifications were filtered according to a 1% false discovery rate at both the peptide and protein group levels. The MaxLFQ method was used for protein quantification and normalization using the total ion current was applied.

#### Library search

DDA raw files from oocyte fractionation were processed using PEAKS Studio 13.1. Data were searched against the SwissProt Mus musculus (01 2026, 17247 entries) database. Semi-specific tryptic cleavages were selected and a maximum of 2 missed cleavages were allowed. The following post-translational modifications were considered for identification: Acetyl (Protein N-term), Oxidation (M), Deamidation (NQ), Carbamylation (K, N-term), Formylation (K, N-term), Phosphorylation (S, T, Y), Methylation (K, R), Dimethylation (K, R), Acetylation (K), Ubiquitin-K (K) as variable and Beta-methylthiolation (C) as fixed. The maximum number of variable modifications per peptide was set to 2. Identifications were filtered according to a 1% false discovery rate at both the peptide and protein group levels.

#### PTM search

DIA raw files were processed with PEAKS Online 13 build 2.2 using a combined library and database search. Search parameters were identical to those used for the library search except that the maximum number of variable modifications per peptide that was set to 2. Identifications were filtered according to a 1% false discovery rate at both the peptide and protein group levels and peptides were quantified using the embedded LFQ quantification module of the software.

#### Statistics

Multivariate statistics on protein measurements were performed using Qlucore Omics Explorer 3.9 (Qlucore AB, Lund, *SWEDEN*). A positive threshold value of 1 was set to allow a log2 transformation of abundance data for normalization *i.e.* all abundance data values below the threshold are replaced by 1 before transformation. The transformed data were finally used for statistical analysis *i.e.* the evaluation of differentially present proteins between two groups using a bilateral Student’s t-test.

## Acknowledgements

We thank team members for discussion and critical reading of the manuscript. We are grateful to Damien Cladière for genotyping and mouse husbandry. We thank administrative and informatics services at the Institut Jacques Monod and members of the animal house “Animalerie Buffon”. KW and SAT received funding for this work from the Agence Nationale de la Recherche (ANR-23-CE13-0015-01 and ANR-21-CE13-0026, respectively). Work in the Wassmann lab was additionally supported by the Fondation pour la Recherche Médical (Equipe FRM DEQ 202103012574).

---

**Figure S1.**
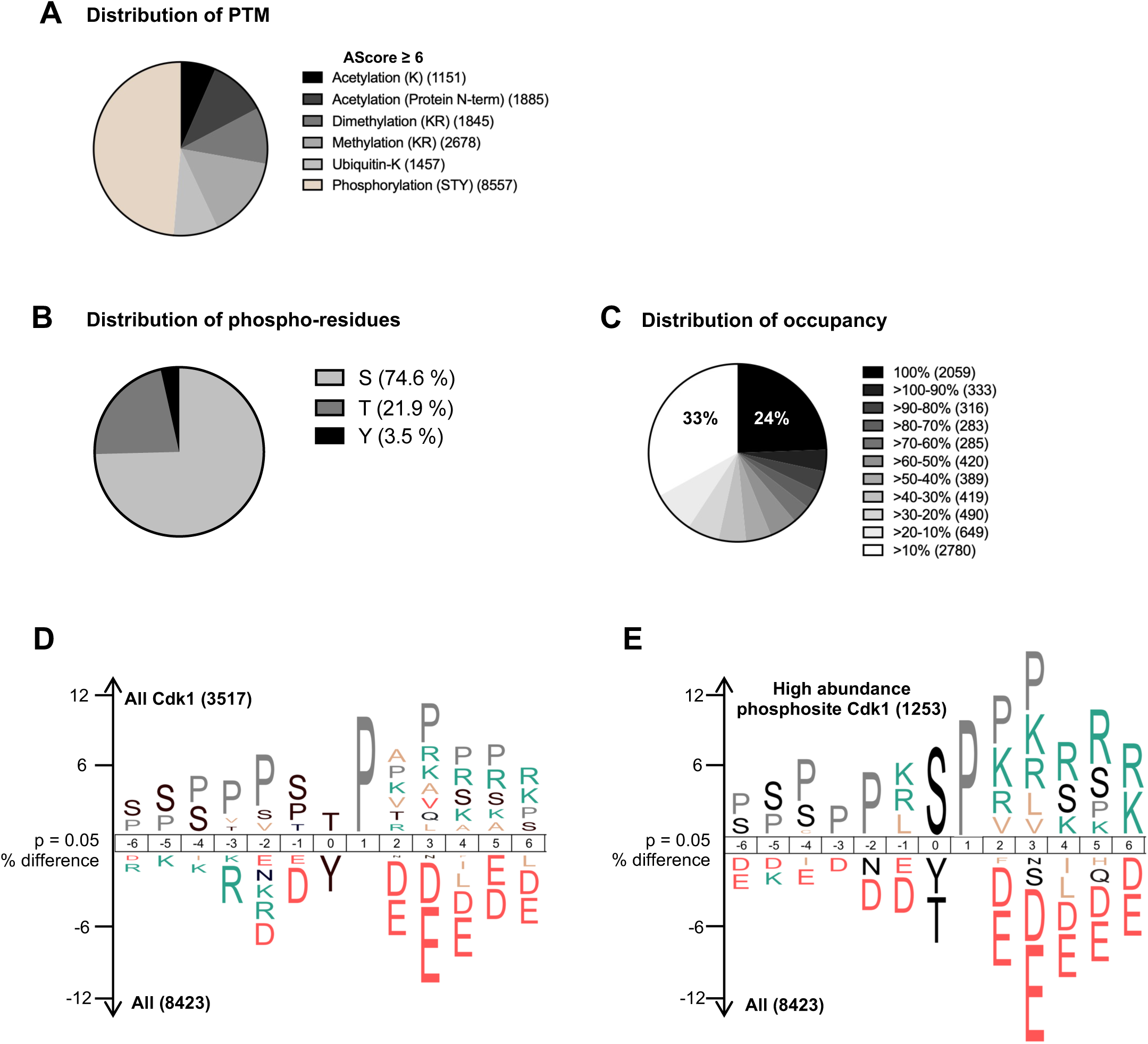
Global features of PTMs, phosphosite composition, phospho-abundance, and motif enrichment. (A) Distribution of the PTMs with AScore ≥ 6. (B) Distribution of phosphorylated residues across the phosphoproteome, showing the percentage of serine (S), threonine (T), and tyrosine (Y) residues (pie chart). (C) Distribution of phosphosites across different phospho-abundance categories, with the number of phosphosites in each category indicated in parentheses (pie chart). (D–E) IceLogo motif analysis (p = 0.05) showing enrichment of sequence motifs. (D) Comparison of Cdk1 phosphosites versus the remainder of the phosphoproteome. (E) Comparison of Cdk1 phosphosites with 100% phospho-abundance versus the remainder of the phosphoproteome. Proline residues at the +1 position were downscaled in the sequence logos for improved visualization.

**Figure S2.**
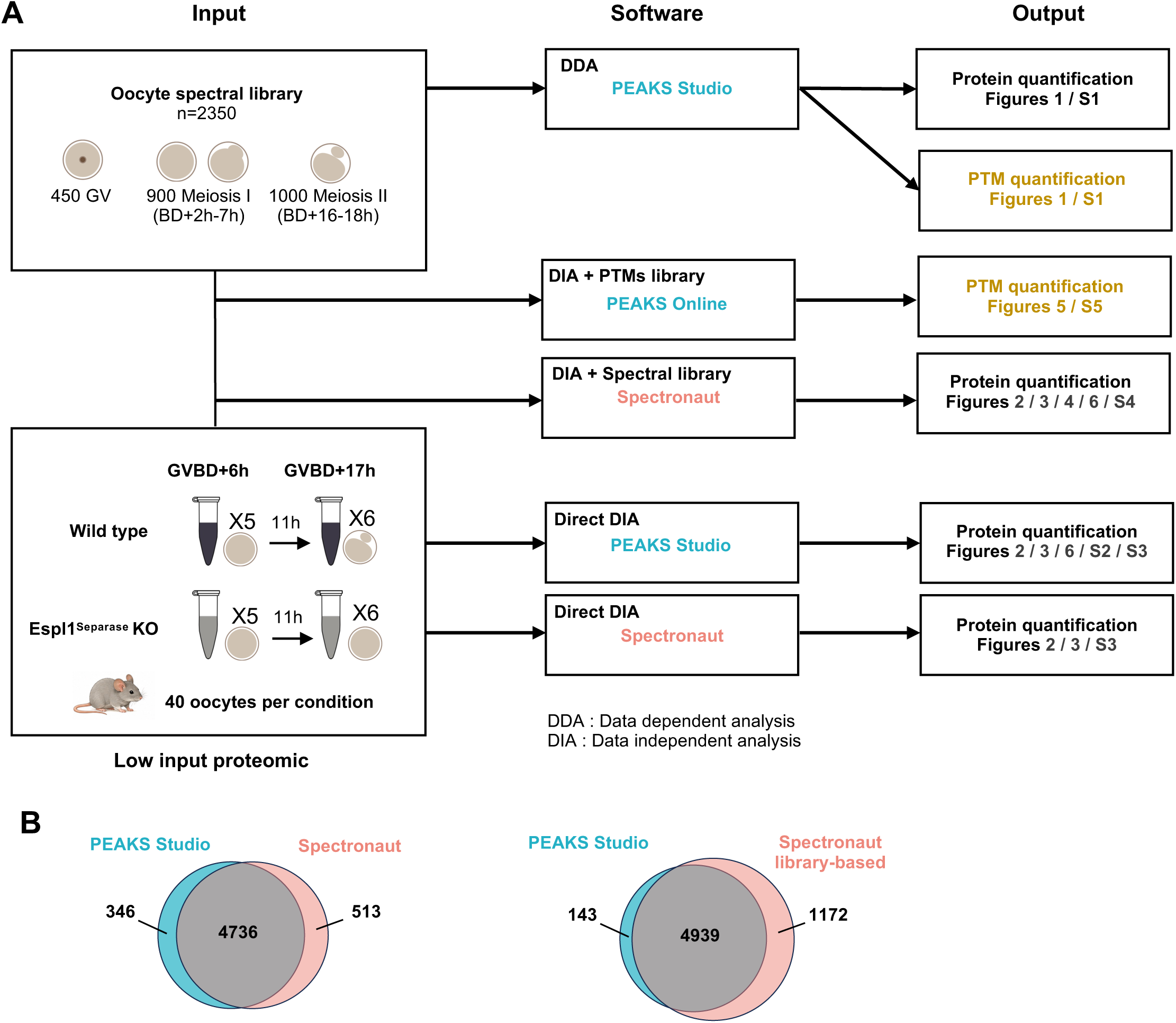
Overview of experimental workflow and data processing pipeline. (A) Schematic representation of the experimental workflow, from sample input to processed data. Software used for analysis and, when applicable, utilization of the spectral library is indicated. Data acquisition mode (DDA or DIA) is shown for each dataset. The type of output and the corresponding figure panels where the data are further analyzed are indicated. (B) Venn diagram showing the overlap between proteins quantified in PEAKS Studio and in Spectronaut using (right) or not (left) the spectral library, illustrating the reproducibility and complementarity of these two DIA-based analysis approaches.

**Figure S3.**
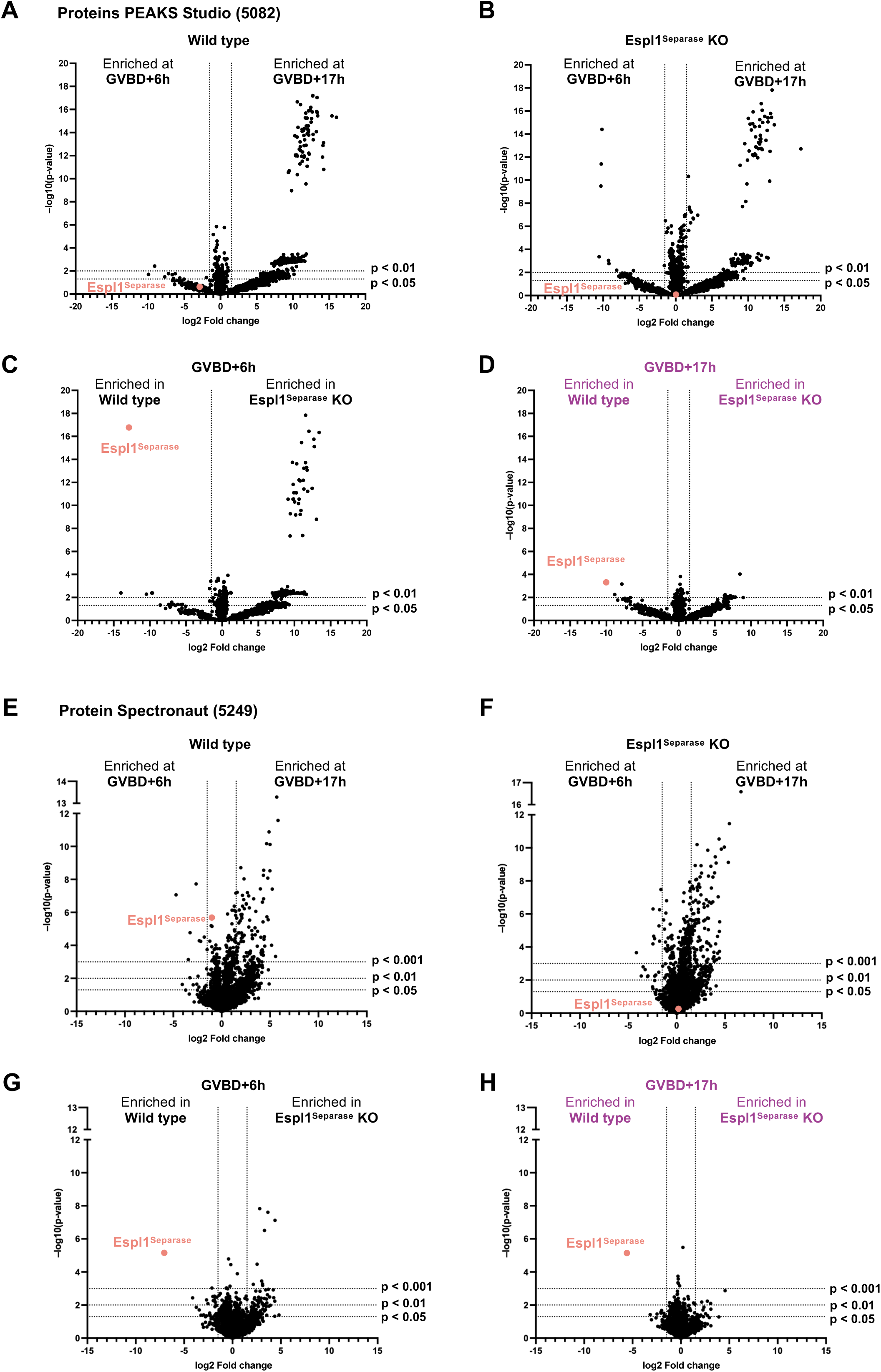
Comparison of differential analysis using PEAKS Studio and Spectronaut (library-free) (A-D) Volcano plots for the four pairwise comparisons: WT GVBD+6h vs WT GVBD+17h, Espl1^Separase^ KO GVBD+6h vs Espl1^Separase^ KO GVBD+17h, WT vs Espl1^Separase^ KO at GVBD+6h, and WT vs Espl1^Separase^ KO at GVBD+17h. Dashed lines indicate the significance thresholds (fold-change ≥ 1.5 combined with either p < 0.05, p < 0.01 or p < 0.001, as indicated). The Espl1^Separase^ protein is highlighted in red in all plots and the WT and Espl1^Separase^ KO mouse at GVBD+17h is highlighted in purple. Data were processed on PEAKS Studio software. (E-H) Same as A-D but data were processed on Spectronaut software.

**Figure S4.**
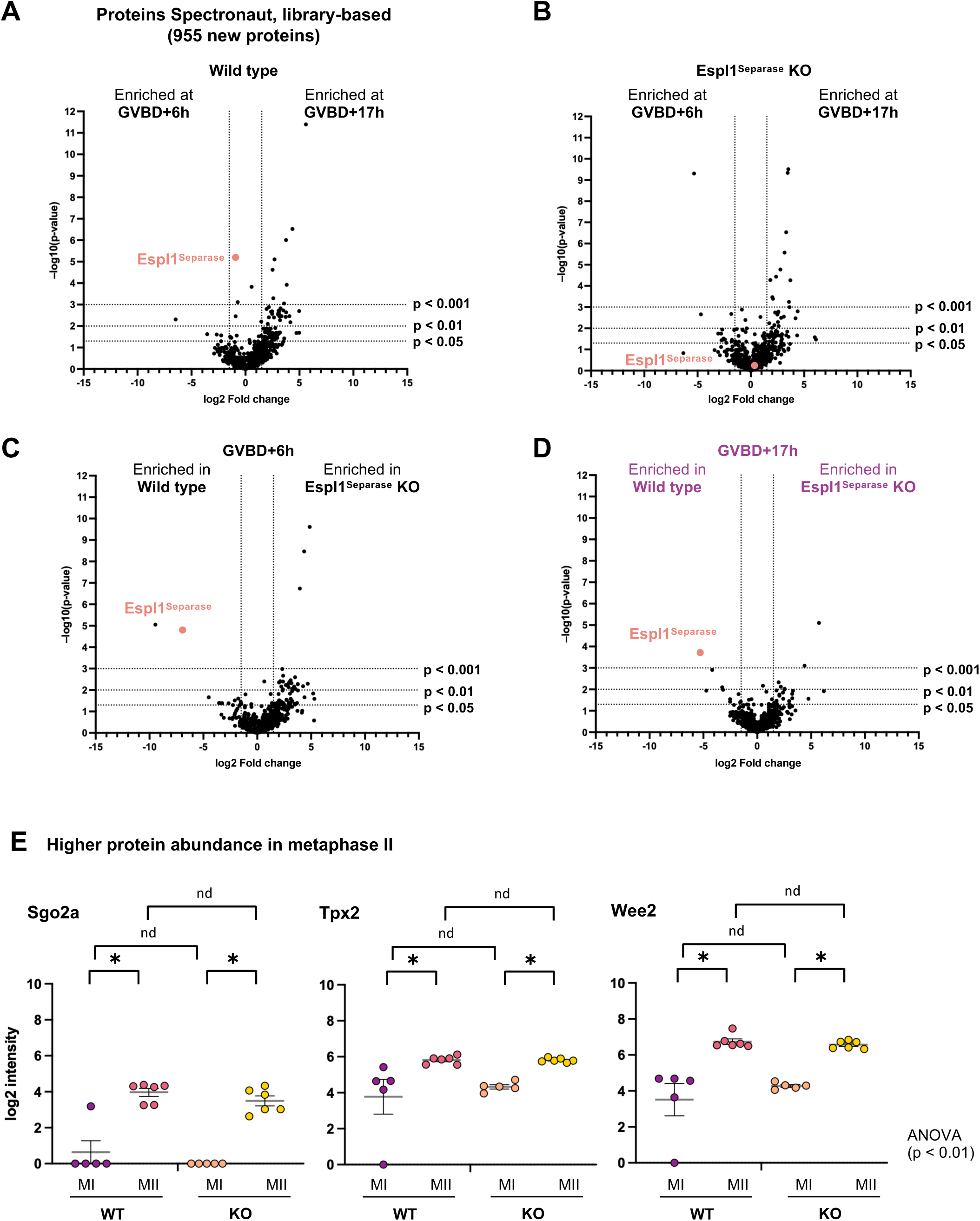
Extra analysis using Spectronaut and spectral library-based approaches. (A-D) Volcano plots for the four pairwise comparisons: WT GVBD+6h vs WT GVBD+17h, Espl1^Separase^ KO GVBD+6h vs Espl1^Separase^ KO GVBD+17h, WT vs Espl1^Separase^ KO at GVBD+6h, and WT vs Espl1^Separase^ KO at GVBD+17h. Dashed lines indicate the significance thresholds (fold-change ≥ 1.5 combined with either p < 0.05, p < 0.01 or p < 0.001, as indicated). The Espl1^Separase^ protein is highlighted in red in all plots and the WT and Espl1^Separase^ KO mouse at GVBD+17h is highlighted in purple. Data were processed with Spectronaut using the spectral library. Only proteins newly identified compared to Spectronaut without the library are shown. (E) Plots of log2-transformed protein intensities for four conditions: WT MI, WT MII, KO MI, and KO MII (n = 5-6 biological replicates per condition). Each point represents a single protein measurement. One-way ANOVA was used to assess global variation (p-value is indicated directly on the figure), followed by FDR-corrected pairwise post-hoc t-tests. Statistical significance after FDR correction (0.05) is indicated by asterisks *. ‘nd’ indicates proteins that did not reach significance. Asterisks denote “discovery”, not the exact q value. Boxplot indicates Mean with SEM.

**Figure S5.**
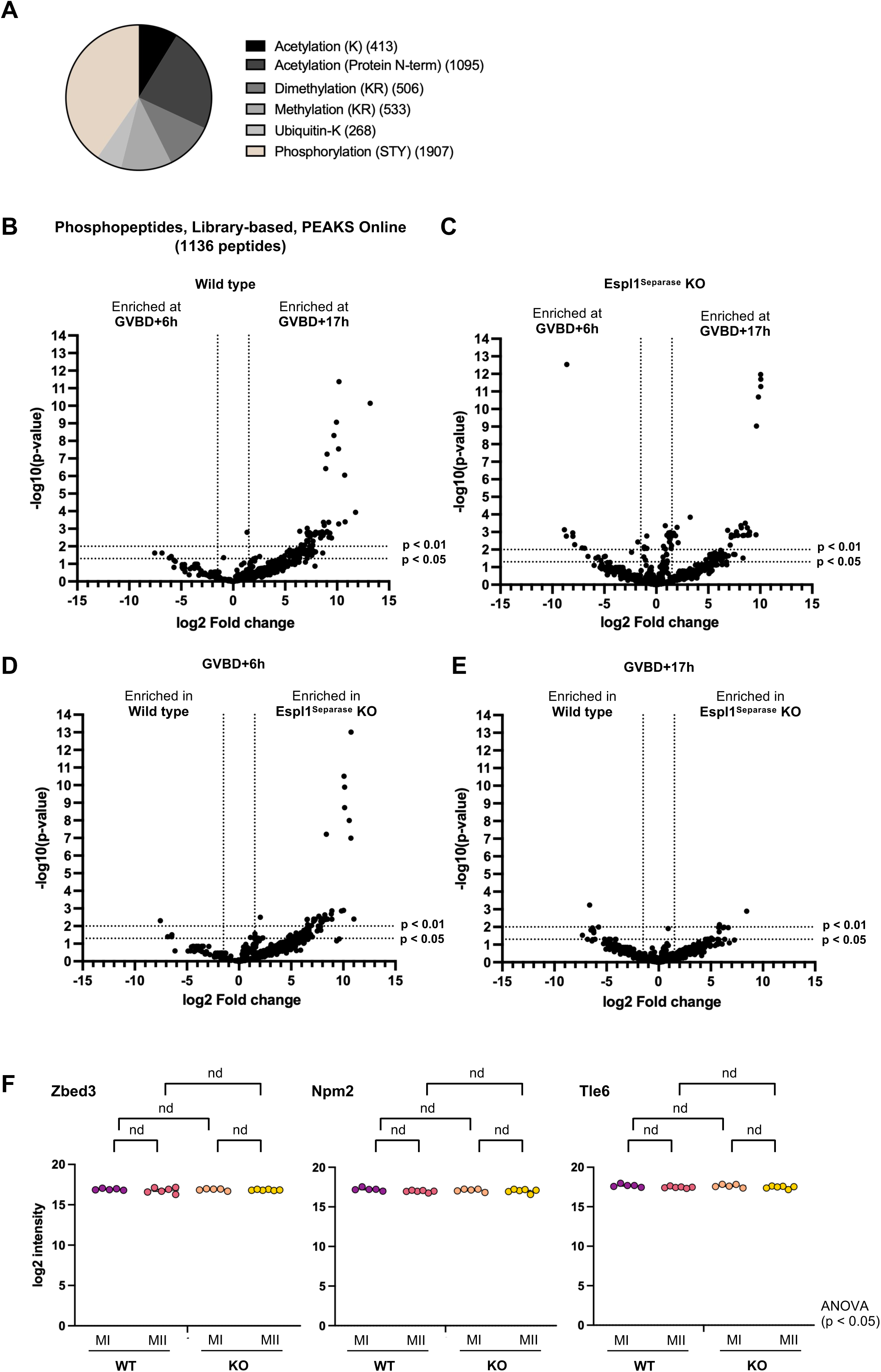
Extra analysis of peptides with PTMs and single phosphosites. (A) Repartition of PTMs categories identified in the proteome and phosphoproteome. (B-E) Volcano plots for the four pairwise comparisons: WT GVBD+6h vs WT GVBD+17h, Espl1^Separase^ KO GVBD+6h vs Espl1^Separase^ KO GVBD+17h, WT vs Espl1^Separase^ KO at GVBD+6h, and WT vs Espl1^Separase^ KO at GVBD+17h. Dashed lines indicate the significance thresholds (fold-change ≥ 1.5 combined with either p < 0.05 or p < 0.01, as indicated). (F) Plots of log2-transformed protein intensities for four conditions: WT MI, WT MII, KO MI, and KO MII (n = 5-6 biological replicates per condition). Each point represents a single protein measurement. One-way ANOVA was used to assess global variation (q or p-value is indicated directly on the figure), followed by FDR-corrected pairwise post-hoc t-tests. Statistical significance after FDR correction (0.05) is indicated by asterisks *. ‘nd’ indicates proteins that did not reach significance. Asterisks denote “discovery”, not the exact q value.

